# A field-ready molecular workflow for sample-to-result detection of the harmful dinoflagellate species *Prorocentrum cordatum* in coastal waters

**DOI:** 10.64898/2025.12.08.692977

**Authors:** Andre Akira Gonzaga Yoshikawa, Beatriz Villefort Rocha, João Victor Costa Guesser, Sabrina Fernandes Cardoso, Iara Carolini Pinheiro, Carlos Yure Barbosa Oliveira, Paulo Antunes Horta, Leonardo Rubi Rorig, Luísa Damazio Pitaluga Rona, André Nobrega Pitaluga

## Abstract

The globally distributed dinoflagellate *Prorocentrum cordatum* contributes to harmful algal blooms, posing serious risks to marine ecosystems and coastal economies. Conventional monitoring of toxic phytoplankton is time-consuming, requires large sample volumes, and depends on expert taxonomic identification, highlighting the need for faster, more sensitive and widely accessible methods. This study presents a sample-to-result molecular assay for detection *P. cordatum* using polymerase chain reaction (PCR) and loop-mediated isothermal amplification (LAMP) targeting the ITS and *psbA* genes. Analyses were performed on small volumes of water (5-50 mL), concentrated with syringe filters and membranes. Sensitivity was evaluated through 10-fold serial dilutions of *P. cordatum* DNA, and specificity tested against three other potentially harmful dinoflagellates, *P*. *lima*, *P*. *hoffmannianum*, and *Takayama acrotrocha*. Detection limits were 5.3 × 10^2^ cells L^−1^ for ITS-PCR and 5.3 × 10^3^ cells L^−1^ for *psbA*-PCR; and 5.3 × 10 and 5.3 × 10 cells L^−1^ for ITS-LAMP and *psbA*-LAMP, respectively. In field tests (n = 48, all < 10^6^ cells L^−1^), *P. cordatum* was detected in 32 samples by ITS-PCR and 29 by *psbA*-PCR. By contrast, ITS-LAMP detected the species in only three samples, and *psbA*-LAMP in none. Overall, PCR proved highly sensitive and specific for detecting low cell densities, while LAMP, although less sensitive, offers a rapid and portable option for field monitoring during moderate-to-high bloom conditions.

**Highlights:** • *P. cordatum* was isolated for molecular assays development.

• Biomass concentration was achieved using small volumes.

• PCR and LAMP assays targeting ITS and *psbA* were validated.

## INTRODUCTION

*Prorocentrum cordatum* (Dinophyceae, syn. *Prorocentrum minimum*) is a bloom-forming marine microalga with a cosmopolitan distribution and is listed in the Taxonomic Reference List of Harmful Microalgae (Lundholm *et al*., 2009; Heil *et al*., 2005). In Brazil, *P. cordatum* has been reported mainly in southern states, occurring year-round in Paranaguá, Paraná, where it has been detected at high cell densities (Mafra Jr. *et al*., 2006; Mafra Jr. *et al*., 2024). In Santa Catarina, records include Penha (2007; Miotto and Tamanaha, 2012) and a mollusk farm in Florianópolis (2006; Alves *et al*., 2010).

Although cases of human intoxication linked to *P. cordatum* have been reported, its toxicity and associated toxins remain uncertain, partly due to possible interactions with bacteria and biphasic blooms involving *Dinophysis acuminata* (Grzebyk *et al*., 1997; Heil *et al*., 2005). This species has multiple nutrient uptake strategies and a mixotrophic metabolism, allowing it to thrive in nitrogen- and phosphorus-enriched environments, including hypoxic regions (Henrichs and Clegg 2024), and to expand geographically in response to human activities (Stoecker *et al*., 1997; Glibert *et al*., 2008; Heil *et al*., 2005; Khanaychenko *et al*., 2019). Moreover, *P. cordatum* belongs to a group of organisms that benefit from coastal pollution, climate change, and ocean acidification (Findlay *et al*. 2025), which increase its photochemical efficiency, energy flux, and antioxidant capacity (Zhu *et al*. 2024). Consequently, *P. cordatum* is considered a sentinel species of eutrophication in coastal ecosystems under the current climate emergency.

Traditional monitoring of harmful microalgae relies mainly on morphological identification and cell counts by light microscopy, which limits scalability, precision, and timely detection. These constraints have driven the development of molecular diagnostic approaches. Among them, Polymerase Chain Reaction (PCR) and Loop-Mediated Isothermal Amplification (LAMP) are highly sensitive and specific tools for identifying dinoflagellates (Toldrà *et al*., 2020). Quantitative PCR (qPCR) and amplicon sequencing have effectively detected *P*. *cordatum* in eastern Australian waters, advancing research on harmful microalgae (McLennan *et al*., 2021). However, PCR requires fully equipped laboratories and trained personnel, restricting its use in the field. By contrast, LAMP, which relies on isothermal amplification, requires minimal equipment and is well suited for on-site monitoring (Toldrà *et al*., 2020). LAMP assays have already been developed for several harmful algal species. For example, Zhang *et al*. (2014) targeted the large subunit (LSU) of rDNA to detect *P. cordatum* using a rapid, low-cost, heat-based DNA extraction method from centrifuged biomass. Similarly, Lee *et al*. (2021) designed LAMP assays targeting the internal transcribed spacer (ITS) and LSU regions for *Ostreopsis* cf. *ovata* and *Amphidinium massartii*, using syringe filters to concentrate biomass from water samples. Together, these studies highlight the potential of LAMP as a practical molecular tool for the field-based monitoring of harmful algal blooms.

Selecting an appropriate molecular marker is critical for nucleic acid amplification assays. The ITS region is widely used to identify harmful algal species due to its high variability (Granéli and Turner, 2006) and has been successfully applied in both PCR and LAMP assays for detecting various dinoflagellates, including *P. cordatum* (Huang *et al*., 2017; Huang *et al*., 2020; Lee *et al*., 2021; McLennan *et al*., 2021; Qin *et al*., 2019; Wang *et al*., 2019; Yang *et al*., 2024; Zhang *et al*., 2012). Plastid genes also hold potential for identifying phototrophic dinoflagellate, but this remains less explored. Among them, the D1 region of the *psbA* gene in chloroplast DNA (cpDNA) has shown promise, revealing phylogenetic clustering of monophyletic species groups at the amino acid level (Granéli and Turner, 2006; Takishita and Uchida, 1999).

This study aimed to develop a sample-to-result molecular approach for monitoring *P*. *cordatum* and to test it in a restricted coastal lagoon on the eastern side of Santa Catarina Island, southern Brazil. Biomass was concentrated from small water volumes (5-50 mL) using a syringe filter and membrane-based protocol. Primers targeting the ITS and *psbA* genes were designed for both PCR and LAMP assays, with sensitivity and specificity assessed. The assays were validated using natural and spiked field samples, and PCR amplicons were sequenced and analyzed phylogenetically to confirm species identity in environmental samples.

## METHODS AND MATERIALS

### Sample collection and algal culture

Four potentially toxic dinoflagellate strains were used in this study: *P. cordatum, Prorocentrum lima, Prorocentrum hoffmannianum*, and *Takayama acrotrocha*. *Prorocentrum cordatum* was collected with a phytoplankton net from Conceição lagoon, a coastal lagoon in Florianópolis, Brazil (Figure 1). The lagoon ranges from mesotrophic to hypertrophic conditions (Silva *et al*., 2017), favouring harmful algal blooms. *Prorocentrum cordatum* (Figure 2) was identified following Steidinger (1997) and isolated using the single-cell capillarity method (Andersen, 2004). The other species were obtained from the Laboratório de Ficologia (LAFIC-UFSC) and the Laboratório de Microalgas (CEM-UFPR).

**Figure 1.**
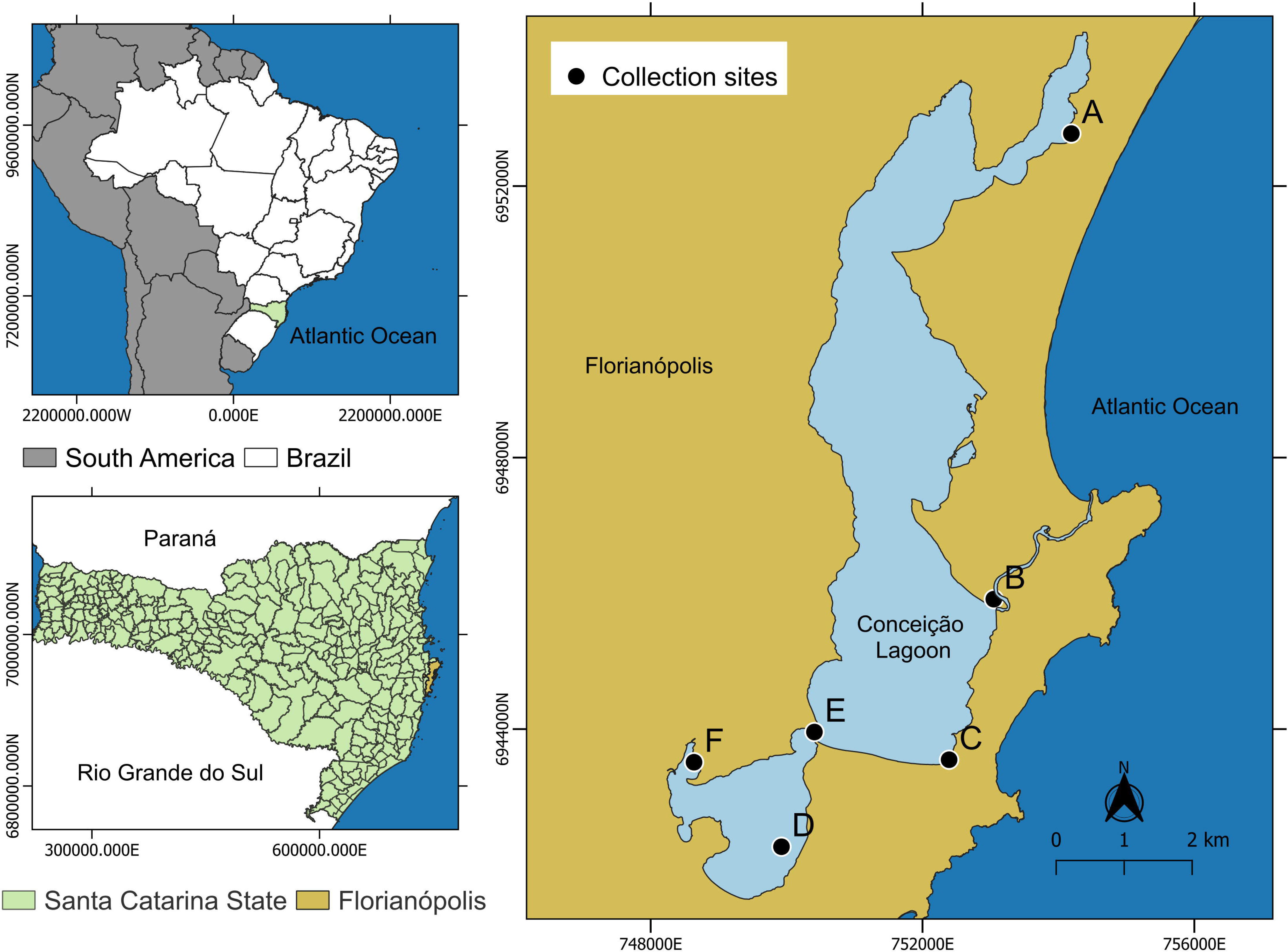
Sampling sites for *Prorocentrum cordatum* monitoring in Conceição lagoon, Florianópolis - SC, Brazil. The top-left inset shows a map of Brazil with the state of Santa Catarina (SC) highlighted in green. Below, a zoomed-in view of SC highlights the municipality of Florianópolis in brown. To the right, a detailed map of Florianópolis shows local water bodies in blue and sampling sites labelled alphabetically. The map’s x- and y-axes represent UTM coordinates. Maps were generated using Quantum GIS (QGIS, version 3.34; QGIS Development Team, 2023).

**Figure 2.**
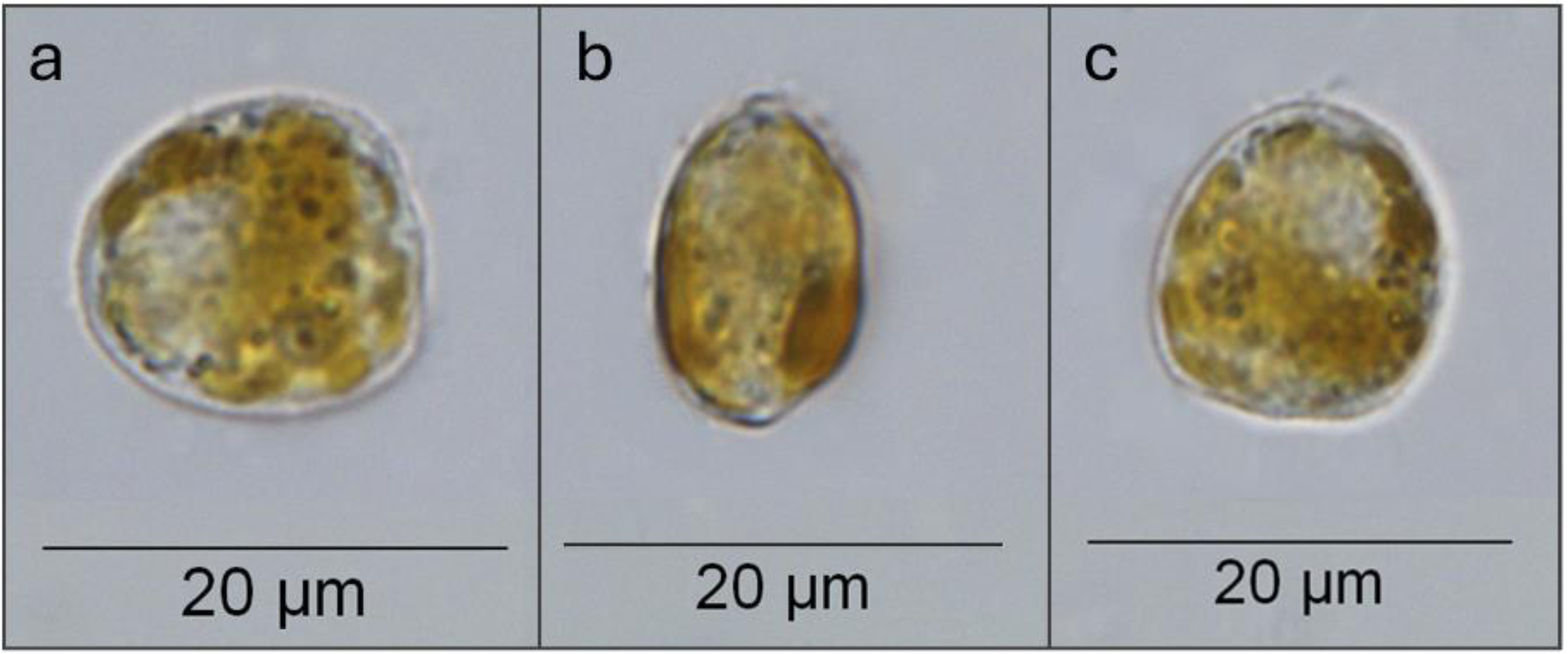
*Prorocentrum cordatum* observed under light microscopy using an Olympus BX-41 microscope at 100× magnification, with measurements performed in ImageJ. (a) Dorsal view of a cell, length = 15.229 μm; (b) dorsal view of an elongated cell, width = 9.251 μm; (c) ventral view of a cell, width = 13.742 μm.

*Prorocentrum cordatum* and *T. acrotrocha* were cultured in 75 mL of modified f/2-Si medium, while *P*. *lima*, and *P*. *hoffmannianum* were maintained in modified f/4-Si medium (Guillard, 1975). All cultures were grown at 20-22°C, salinity 30, under ∼100 μmol photons m^-2^ s^-1^ with a 12:12 h light-dark cycle and subcultured every two months. For cell counts, exponential-phase cultures were gently homogenized, fixed with 1% acid Lugol solution, and counted in triplicate using a Neubauer chamber under a light microscope.

### Biomass concentration method and DNA extraction

Exponential-phase cultures (see *Sample collection and algal culture*) and field samples (see *Field sample validation*) were homogenized and concentrated using syringes (Descarpack), syringe filters (Millipore SX0002500 and SX0001300), and membranes (Millipore) (Supplementary Figure 1). The biomass concentration protocol was adapted from Lee *et al*. (2021). For field samples, a 100 μm nylon pre-filter (Millipore NY1H2500) was used to remove macroalgae, zooplankton, and other large particles (Supplementary Figure 1.b). Subsequently, 10-60 mL of culture or field sample were drawn into syringes and filtered through a 0.22 μm polyether sulfone (Millipore GPWP02500) or mixed cellulose ester membrane (Millipore GSWP01300) to concentrate biomass (Supplementary Figure 1.c). The biomass-containing filter was then rinsed with 2 mL of nuclease-free water to remove salts and dissolved substances. Membranes were dried on a paper towel for two minutes and stored at -80 °C until further analysis (Supplementary Figure 1.d).

Genomic DNA was extracted from retained biomass using either the DNeasy Blood & Tissue Kit (Qiagen), following adaptations from Eland *et al*. (2012), or a heat-based method adapted from Zhang *et al*. (2014). In the latter, membranes were eluted with 100 μL nuclease-free water and heated at 100 °C for 10 minutes. DNA concentration was quantified using the Qubit dsDNA BR Assay Kit (ThermoFisher Scientific, Q32851) on a Qubit 4 Fluorometer. All DNA extracts were stored at -20 °C until use.

### PCR primer design and assay

Two sets of PCR primers were developed to detect *P. cordatum*: one targeting the ITS1-5.8S-ITS2 region of nuclear rDNA (PcorITS1 and PcorITS2, Table 1), and another targeting the *psbA* gene of chloroplast cpDNA (PcorpsbAF and PcorpsbAR, Table 1).

**Table 1.**
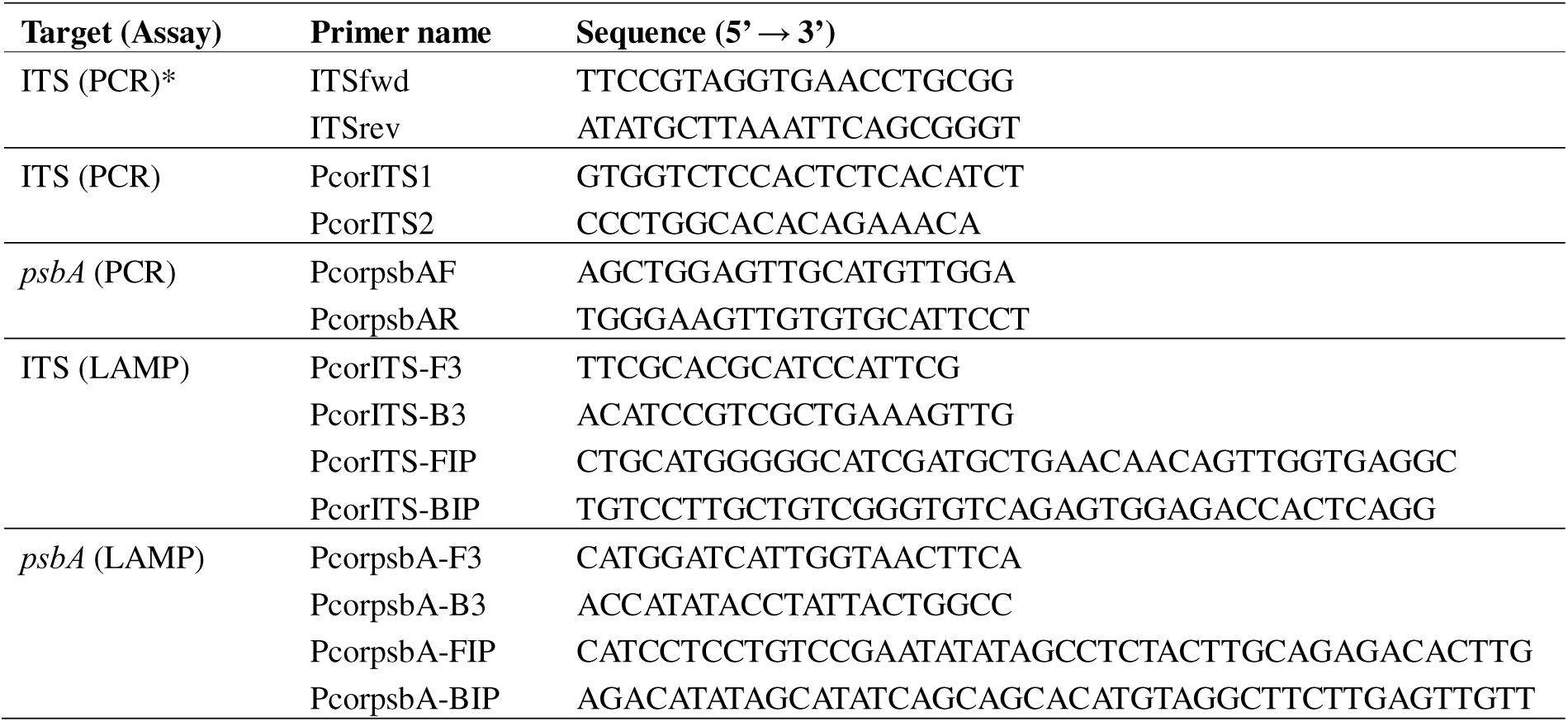
Sequences of primers used in this study. *ITS Primer designed by McLennan *et al*., 2021.

Primers were designed using NCBI Primer-BLAST (Ye *et al*., 2012) with sequences from GenBank (JX402086.1 and JX573313.1) and sequences generated in this study (see *Sequencing and phylogenetic analysis*). Initially, a published primer set (McLennan *et al*., 2021) targeting *P. cordatum* (ITSfwd and ITSrev, Table 1) was used to amplify a 664 bp fragment of the ITS1-5.8S-ITS2 rDNA region from DNA extracted from *P. cordatum* isolated in this study (see *Sample collection and algal culture*). The resulting amplicons (Accession numbers: PV730001-6) were sequenced and, together with the reference sequence JX402086.1, served as templates for designing new primers (PcorITS1 and PcorITS2, Table 1) targeting a shorter 243 bp region. The *psbA* primers (PcorpsbAF and PcorpsbAR, Table 1) were designed based on the reference sequence JX573313.1, amplifying a 661bp fragment of the D1 region.

PCR was performed in 25 μL reactions containing 1× buffer, 2 mM MgCl , 0.4 mM dNTPs, 0.4 μM of each primer, 1 U Hot Start Taq polymerase (Promega M7841), 10-20 ng DNA, and nuclease-free water. Amplification was run on a MiniAmp Plus Thermal Cycler (ThermoFisher Scientific) with the following program: initial denaturation at 95 °C for 5 min; 30 cycles of 95 °C for 45 s, annealing at 60 °C (ITS) or 65 °C (*psbA*) for 45 s, and 72 °C for 45 s; final extension at 72 °C for 5 min. Products were visualized by electrophoresis on 1-1.5% agarose gels.

### Sequencing and phylogenetic analysis

PCR amplicons generated with ITSfwd/ITSrev, PcorITS1/PcorITS2, and PcorpsbAF/PcorpsbAR were purified using the Wizard SV Clean-Up System (Promega A9281), following the manufacturer’s instructions, and quantified with a NanoDrop Lite Plus spectrophotometer (ThermoFisher Scientific). Purified fragments were bidirectionally sequenced at the FIOCRUZ Sequencing Unit (RPT1A, Rio de Janeiro, Brazil) on an ABI Prism 3730 DNA sequencer using the ABI Prism Big Dye Terminator Cycle Sequencing kit (Applied Biosystems).

Chromatograms were inspected with CHROMAS version 2.4, and consensus sequences were assembled in SeqMan version 7.0. Species identity was confirmed using BLASTn, and sequences were aligned with ClustalW (Madeira *et al*., 2024). Final sequences were deposited in GenBank (Accession numbers: ITS: PV730001-15; *psbA*: PV785871-83).

Phylogenetic analysis of *P. cordatum psbA* (PV785871-83) and ITS (PV730001-15) sequences from field and culture samples were performed with IQ-TREE v2.4 (Nguyen *et al*., 2015) using the best-fit models TVM+F+I and K2P+G4, respectively. Trees were visualized with iTOL (Letunic and Bork, 2021).

### Design of LAMP primers and assay Implementation

Consensus ITS1-5.8S-ITS2 (PV730001-6) and *psbA* (PV785877-80) sequences generated in this study were aligned with JX402086.1 and JX573313.1, respectively, using ClustalW (Madeira *et al*., 2024). These alignments were used to design LAMP primers with the NEB LAMP Primer Design Tool v1.4.1 (https://lamp.neb.com/). Primer sets for both regions included a forward primer (F3), backward primer (B3), forward inner primer (FIP), and backward inner primer (BIP) (Notomi *et al*., 2000) (Table 1).

Primers were diluted in nuclease-free water and combined into a 10× primer mix containing 16 μM each of FIP and BIP, and 2 μM each of F3 and B3. LAMP reactions were performed using the WarmStart^®^ Colorimetric LAMP Master Mix (New England BioLabs, M1704) in a final volume of 10 μL, containing 5 μL of 2× Master Mix, 1 μL of 10× primer mix, and 4 μL of template DNA. Reactions were incubated at 65 °C for 60 minutes in 0.2 mL microtubes on a dry bath (Kasvi).

For product detection, 1 μL of 1000× SYBR Green I (Invitrogen S7563) was added to the inner lid of each 0.2 mL tube before incubation and mixed into the reaction after the LAMP reaction was complete. Amplification was detected visually by SYBR Green I through both colour change observable to the naked eye and fluorescence under UV light, and further confirmed by electrophoresis on a 2% agarose gel.

### Specificity and sensitivity analyses of PCR and LAMP assays

For specificity testing, genomic DNA was extracted from four algal isolates, *P*. *cordatum, P*. *lima, P*. *hoffmannianum*, and *T*. *acrotrocha* (see *Sample collection and algal culture*), using the DNeasy Blood & Tissue Kit (Qiagen). The DNA samples were tested using both PCR and LAMP assays, following the conditions described in the sections *PCR primer design and assay* and *Design of LAMP primers and assay implementation*, respectively.

To evaluate sensitivity, genomic DNA from *P. cordatum* (5.3 × 10 cells L^−1^ = 10 ng μL^−1^) was serially diluted ten-fold (10 ng μL^−1^ to 10 ng μL^−1^) and used as template for both PCR and LAMP assays.

To further evaluate LAMP sensitivity in environmental conditions, *P. cordatum* cells in the exponential growth phase were spiked into a water sample from Conceição lagoon (Florianópolis, SC) previously tested for the presence of *P. cordatum* by PCR and Utermöhl analysis. A 4 mL aliquot of the environmental water was inoculated with 1 mL of *P. cordatum* culture (1.2 × 10 cells/L), followed by ten-fold serial dilutions down to 1.2 × 10^2^ cells/L. Each dilution was tested in triplicate using the LAMP assay. Detection limits were estimated using probit regression analysis with MedCalc Statistical Software version 19.2.6 (MedCalc Software bv, Ostend, Belgium; https://www.medcalc.org/; 2020).

Details on the visualization of PCR and LAMP products for both sensitivity and specificity analyses are provided in the sections *PCR primer design and assay* and *Design of LAMP primers and assay implementation*, respectively.

### Assay validation based on field-collected samples

A total of 48 environmental water samples were collected from six sites in Conceição Lagoon (Figure 1; Table 2) during 2024 and 2025 to standardize field analyses. Both concentrated samples (phytoplankton net) and whole water samples (Van Dorn bottle) were included. The samples were grouped according to collection period:(i) January-February 2024, (ii) September 2024, and (iii) January 2025.

**Table 2.**
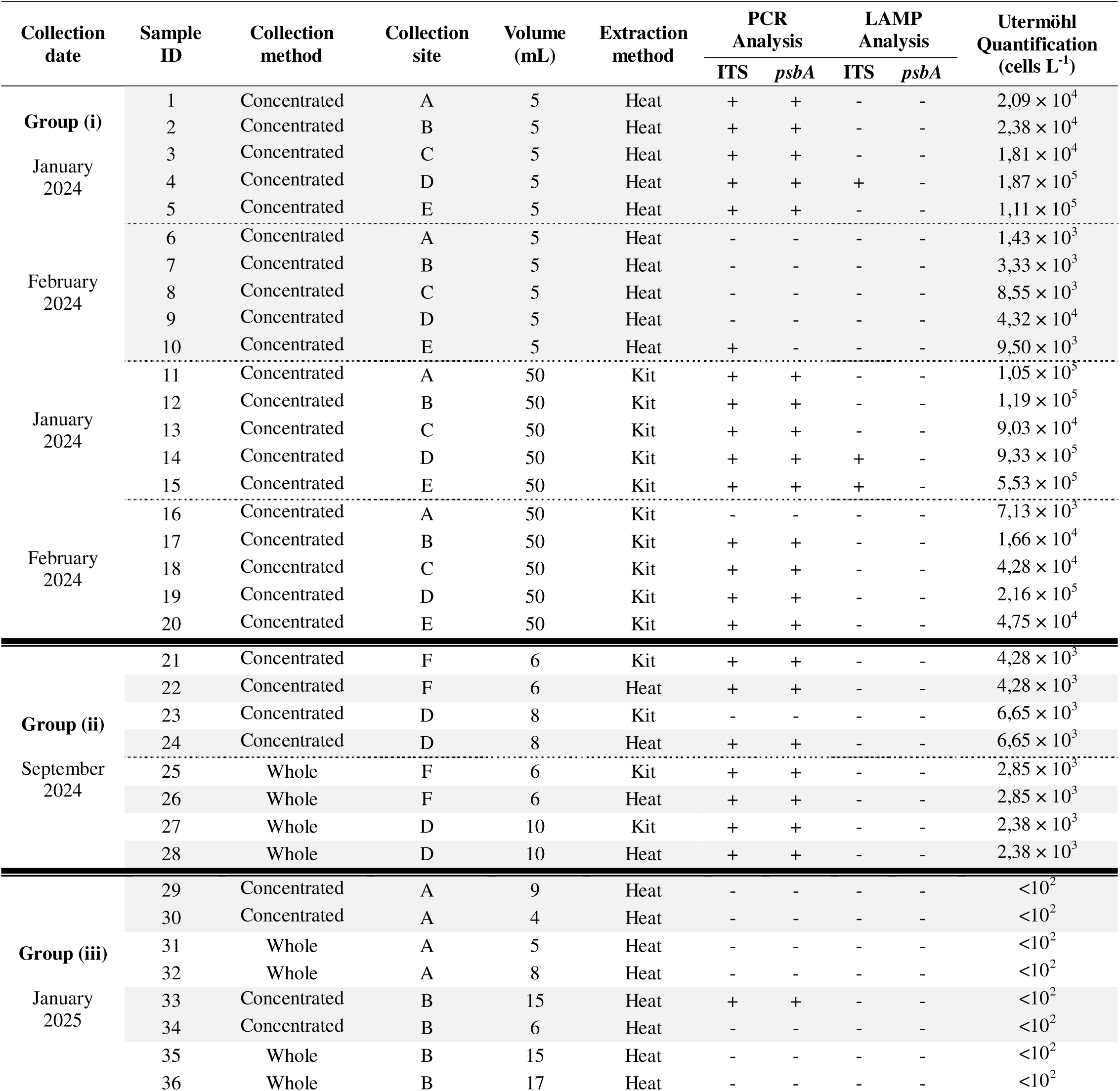
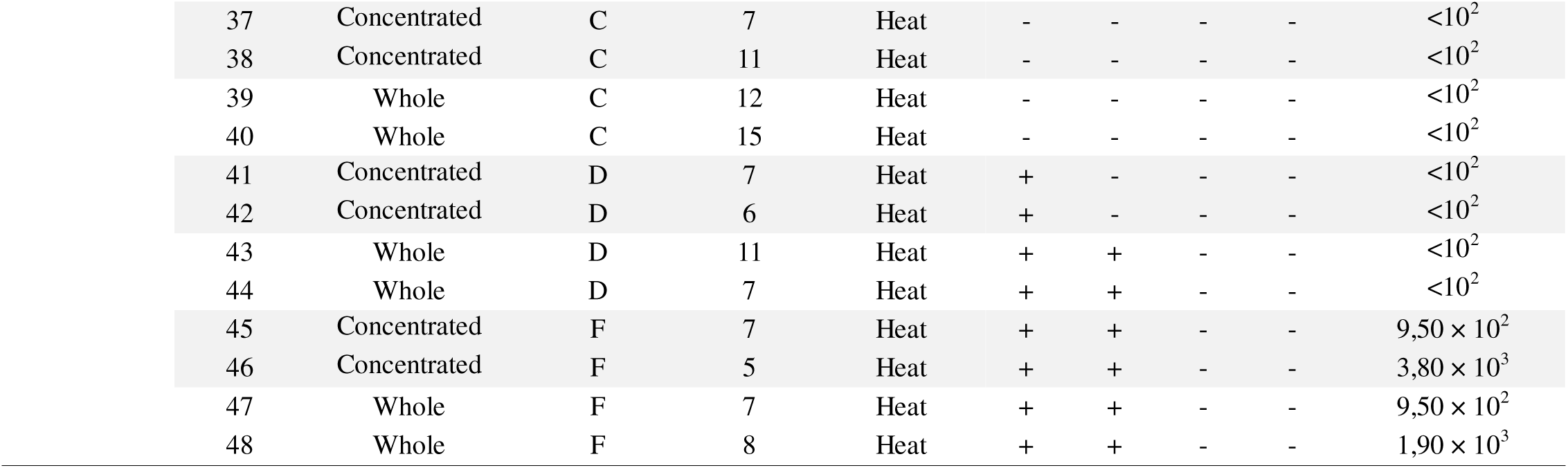
Validation of field samples collected in Conceição Lagoon. Environmental water samples collected from sites A-F in Conceição Lagoon (Figure 1) during 2024 and 2025 and grouped by collection date: Group (i) January-February 2024, Group (ii) September 2024, and Group (iii) January 2025. *Collection method*: concentrated samples collected using a phytoplankton net, or whole water samples collected using a Van Dorn bottle; *Extraction method*: DNA extracted using the heat-based method, or the DNeasy Blood & Tissue Kit (Qiagen). + indicates that amplicons were detected in the assays; – indicates that no amplicons were detected.

Group (i) comprised 20 concentrated samples collected from sites A-E. At each site, 5 mL and 50 mL of water were filtered through 13 mm and 25 mm syringe membranes, respectively. DNA was extracted from the 5 mL samples using a heat-based method, and from the 50 mL samples using the DNeasy Blood & Tissue Kit (Qiagen). Group (ii) included eight samples from sites D and F. Biomass was filtered onto 13 mm membranes until saturation. Concentrated samples saturated at 6 – 8 mL, while whole samples saturated at 6 – 10 mL (Table 2). Biomass was concentrated in duplicate for each collection method, and DNA was extracted using both the heat-based method and the DNeasy Blood & Tissue Kit. Group (iii) consisted of 20 concentrated and whole samples from sites A, B, C, D, and F. As in Group (ii), biomass was filtered onto 13 mm membranes until saturation, with volumes ranging from 4 to 17 mL. DNA was extracted using only the heat-based method.

For cell quantification, each field sample was gently homogenized, and a subsample was fixed with 1% acid Lugol’s solution. Fixed material was transferred to 10 mL Utermöhl chambers (limit of detection: 10^2^ cells L^−1^), settled for 24 hours, and analysed in triplicate using an inverted light microscope (Edler and Elbrächter, 2010).

DNA samples were tested with both PCR and LAMP assays targeting the ITS and *psbA* regions, following the conditions described in the sections *PCR primer design and assay* and *Design of LAMP primers and assay implementation*. Product visualisation methods are also detailed in those sections.

## RESULTS AND DISCUSSION

In this study, *P. cordatum* (Figure 2) was isolated from Conceição lagoon, Brazil, and maintained in laboratory conditions. We developed PCR and LAMP assays targeting the ITS and *psbA* regions and validated them with field samples. The ITS region is widely used in molecular identification due to its high resolution in distinguishing closely related species; for instance, a qPCR ITS-based assay for *P*. *cordatum* has already been established (McLennan *et al*., 2021). Here, we validated a sensitive benchtop PCR assay and a portable LAMP assay capable of detecting abnormal cell concentrations.

In contrast, plastid genes such as *psbA*, although promising for species-level identification, have been less frequently applied in algal taxonomy and molecular diagnostics (Granéli and Turner, 2006). Our study demonstrates the utility of *psbA* as a complementary molecular marker for *P*. *cordatum*, thereby expanding the repertoire of genetic targets for HAB monitoring and offering a foundation for future applications in molecular ecology and environmental surveillance.

### Sensitivity

The sensitivity of the assays was assessed using *P. cordatum* genomic DNA subjected to ten-fold serial dilutions. The ITS-PCR assay detected concentrations as low as 10^4^ ng/μL (∼5.3x10^2^ cells L^−1^), while the *psbA*-PCR detected down to 10^3^ ng/μL (∼5.3 × 10^3^ cells L^1^) (Figure 3). For LAMP, the ITS-based assay detected down to 10 ng/μL (∼5.3 × 10^5^ cells L^1^), whereas the *psbA*-LAMP assay detected only at 10^0^ ng/μL (∼5.3 × 10^6^ cells L^1^) (Figure 4). Overall, ITS assays were ten times more sensitive than *psbA* assays, likely due to differences in gene copy number. The LSU region, which includes the ITS, is estimated at ∼398 ± 184 copies per *P. cordatum* cell (Liu *et al*., 2021), whereas *psbA* copy numbers in dinoflagellates are more variable and generally lower (Barbrook *et al*., 2006, Iida *et al*., 2009). Nevertheless, despite its lower sensitivity, the *psbA*-PCR assay still detected relatively low cell concentrations (5.3 × 10^3^ cells L^1^), highlighting its potential as a low-copy genetic marker. Moreover, PCR was up to 1,000 times more sensitive than LAMP under the tested conditions; however, the reduced sensitivity of LAMP may actually be advantageous for point-of-care surveillance, since dinoflagellates such as *P. cordatum* typically reach hazardous levels only at higher cell densities (Mafra Jr. *et al*., 2024). Therefore, LAMP provides a practical and field-ready tool for routine monitoring.

**Figure 3.**
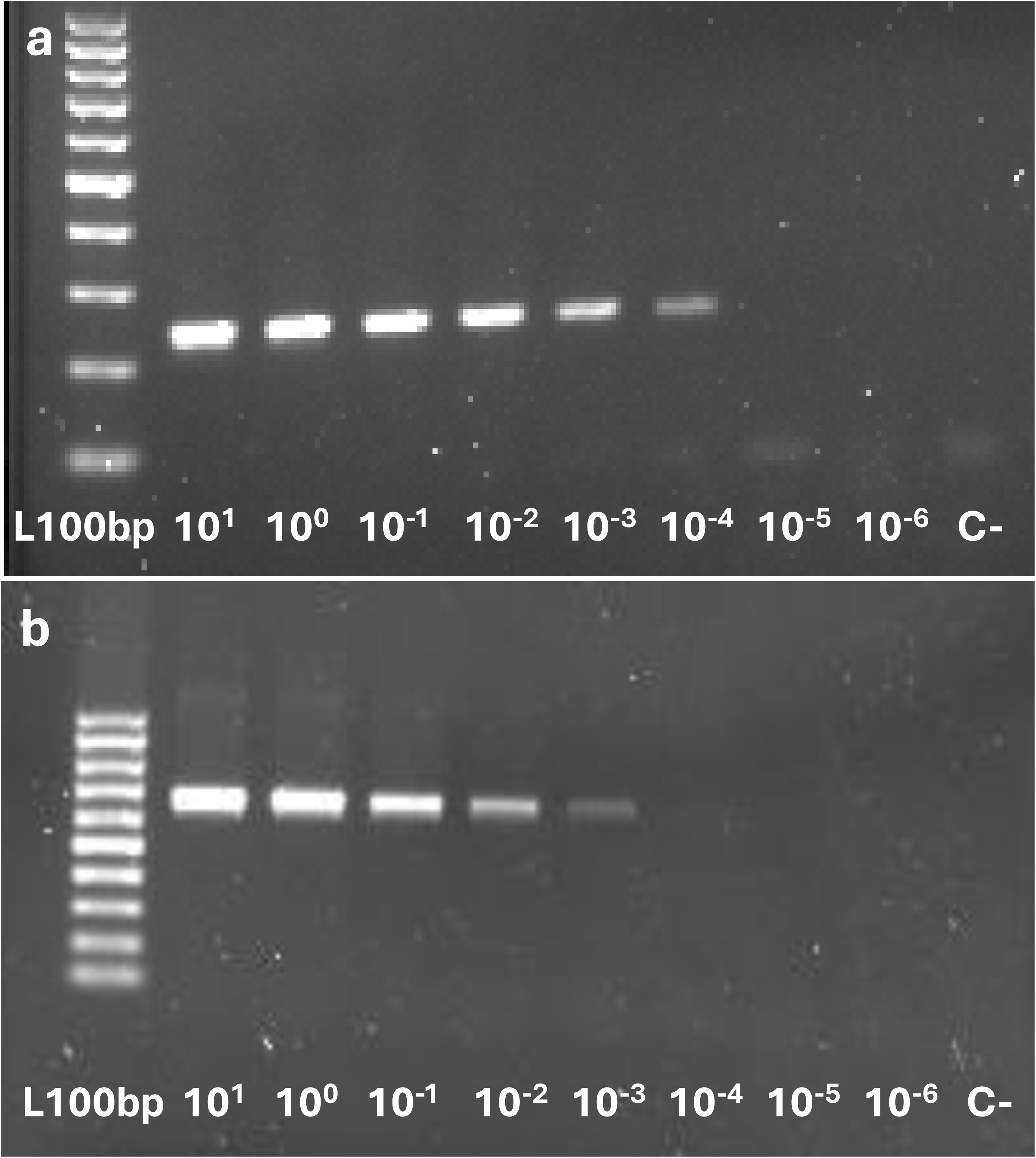
Sensitivity of PCR assays. (a) ITS-PCR (243bp) and (b) *psbA*-PCR (661bp) assays tested with *Prorocentrum cordatum* DNA at concentrations ranging from 10^1^ to 10 ng/μL. Line L100bp: 100 bp molecular marker (100 bp); lines 10^1^ to 10 : serial dilutions of DNA; line C-: negative control (no DNA, water used instead). The ITS-PCR assay demonstrated greater sensitivity than the *psbA*-PCR, detecting DNA concentrations as low as 10 ng/μL (approximately 5.3 × 10^2^ cells L^−1^), while *psbA*-PCR detected down to 10^−3^ ng/μL (∼5.3 × 10^3^ cells L^−1^).

**Figure 4.**
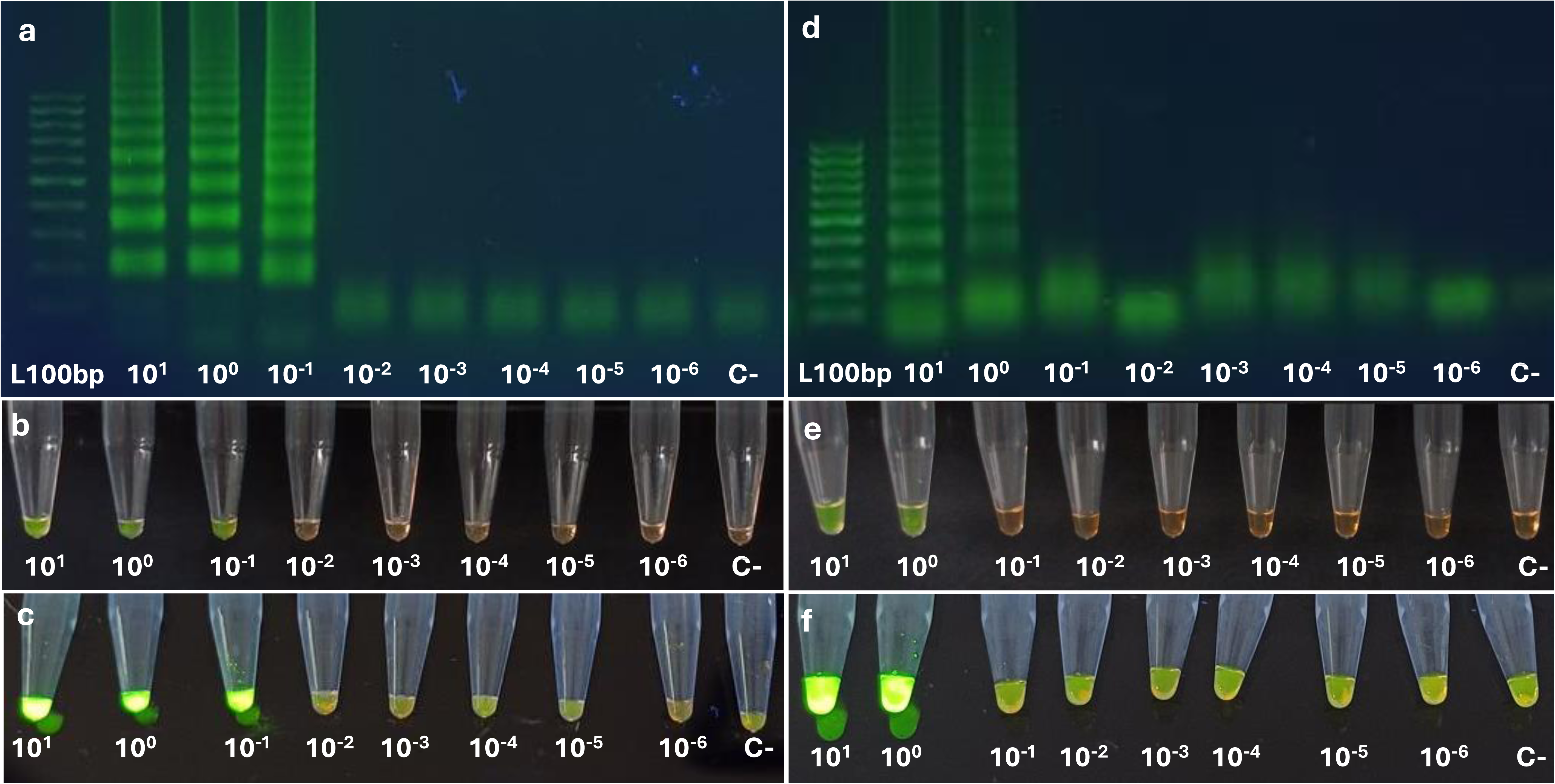
Sensitivity of LAMP assays. Amplification of *Prorocentrum cordatum* DNA using ITS-LAMP and *psbA*-LAMP assays with DNA concentrations ranging from 10^1^ to 10 ng/μL, evaluated by: (a) and (d) electrophoresis on a 2% agarose gel; (b) and (e) visible colour change after staining with 1000 × SYBR Green I (Invitrogen S7563); and (c) and (f) fluorescence under UV light. Panels (a) to (c) show results for the ITS-LAMP assay, and (d) to (f) for the *psbA*-LAMP assay under the same conditions. Line L100bp: 100 bp molecular marker; lanes 10^1^ to 10 : serial DNA dilutions; lane C-: negative control (water instead of DNA). The ITS-LAMP assay showed higher sensitivity, detecting DNA concentrations as low as 10^−1^ ng/μL (approximately 5.3 × 10 cells L^−1^), while the *psbA*-LAMP assay detected only at 10 ng/μL (5.3 × 10 cells L^−1^).

To further assess LAMP assay sensitivity, a water sample from Conceição lagoon was spiked with 1 mL of *P. cordatum* culture, and DNA was extracted using the heat-based method. The sample was serially diluted ten-fold from 1.2 × 10 cells L^−1^ to 1.2 × 10^2^ cells L^−1^ and tested in triplicate with both ITS- and *psbA*-LAMP assays. The ITS-LAMP assay yielded positive results in all triplicates down to 1.2 × 10 cells L^−1^ (Figure 5), with a detection limit of 2.4 × 10^4^ cells L^−1^ (P < 0.0001) estimated by probit regression. For *psbA*-LAMP, all triplicates tested positive at 1.2 × 10 cells L^−1^, with an estimated detection limit of 329,092 cells L^−1^ (P < 0.0001) (Figure 5). These results confirm that LAMP assays effectively detect moderate to high *P. cordatum* cell densities, consistent with the threshold reported by Mafra Jr. *et al*. (2024), who classified the species as potentially harmful above 10,000 cells L^−1^ (intermediate effects) and 100,000 cells L^−1^ (severe effects). Accordingly, our assays were optimised to target cell densities relevant to bloom events, typically exceeding 10 cells L^−1^.

**Figure 5.**
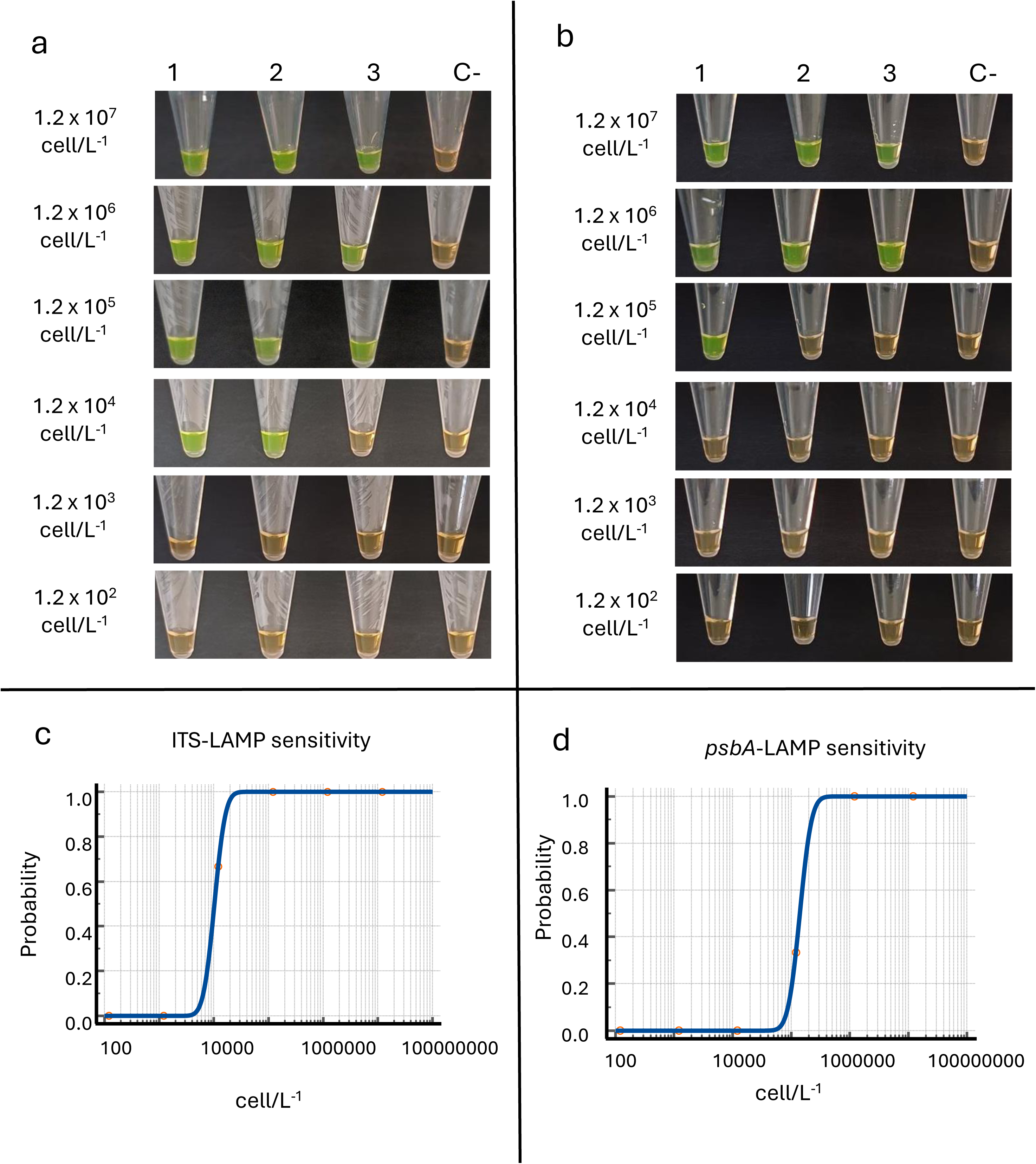
Analytical sensitivity of LAMP assays. Colorimetric detection of *Prorocentrum cordatum* DNA using ITS-LAMP (a) and *psbA*-LAMP (b) assays, evaluated across concentrations from 1.2 × 10 to 1.2 × 10^2^ cells L^−1^, based on visible colour change after staining with 1000× SYBR Green I (Invitrogen S7563). Each assay was run in triplicate (lines 1 to 3), with a negative control (C-, water only). Sensitivity for ITS (c) and *psbA* (d) assays were determined via probit regression. ITS-LAMP showed higher sensitivity, with all triplicates testing positive down to 1.2 × 10 cells L^−1^ and a detection limit of 24,012 cells L^−1^ (P < 0.0001), while the *psbA*-LAMP detected down to 1.2 × 10 cells L^−1^, with a detection limit of 329,092 cells L^−1^ (P < 0.0001).

### Specificity

The specificity of the assays was evaluated using genomic DNA from four algal species: *P. cordatum*, *P. lima*, *P. hoffmannianum*, and *T*. *acrotrocha*. Both PCR (Figure 6) and LAMP (Figure 7) assays targeting the ITS and *psbA* regions yielded specific amplification for *P. cordatum*, with no detectable amplification in the other species.

**Figure 6.**
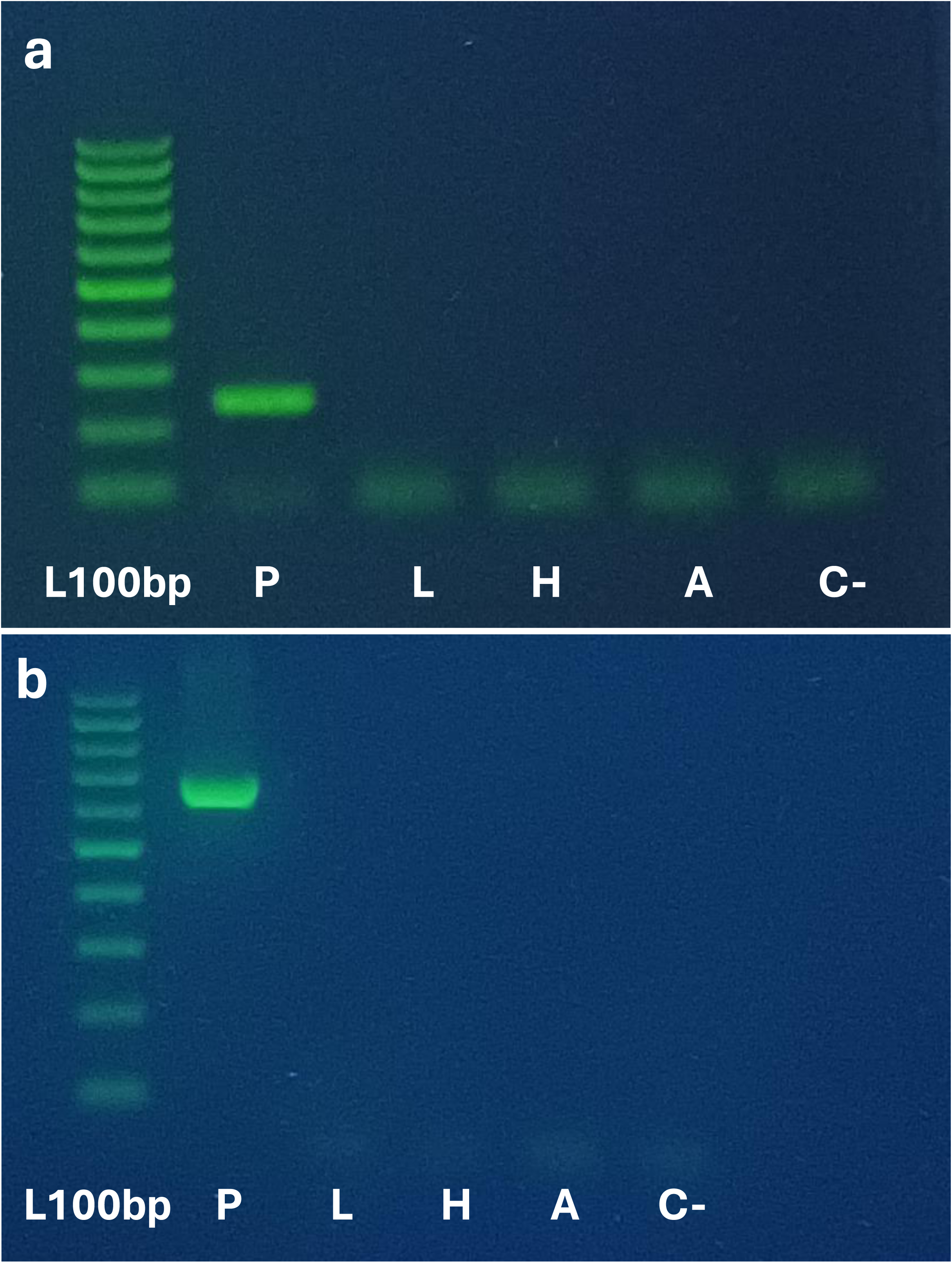
Specificity of PCR assays. (a) ITS-PCR (243bp) and (b) *psbA*-PCR (661bp) assays were tested using DNA from *Prorocentrum cordatum* (P), *Prorocentrum lima* (L), *Prorocentrum hoffmannianum* (H), and *Takayama acrotrocha* (A) DNA at concentrations of 10 ng/μL. Line L100bp: 100 bp molecular marker; line C-: negative control (water, no DNA). Both assays showed 100% specificity for *P. cordatum*, with no amplification observed for the other species.

**Figure 7.**
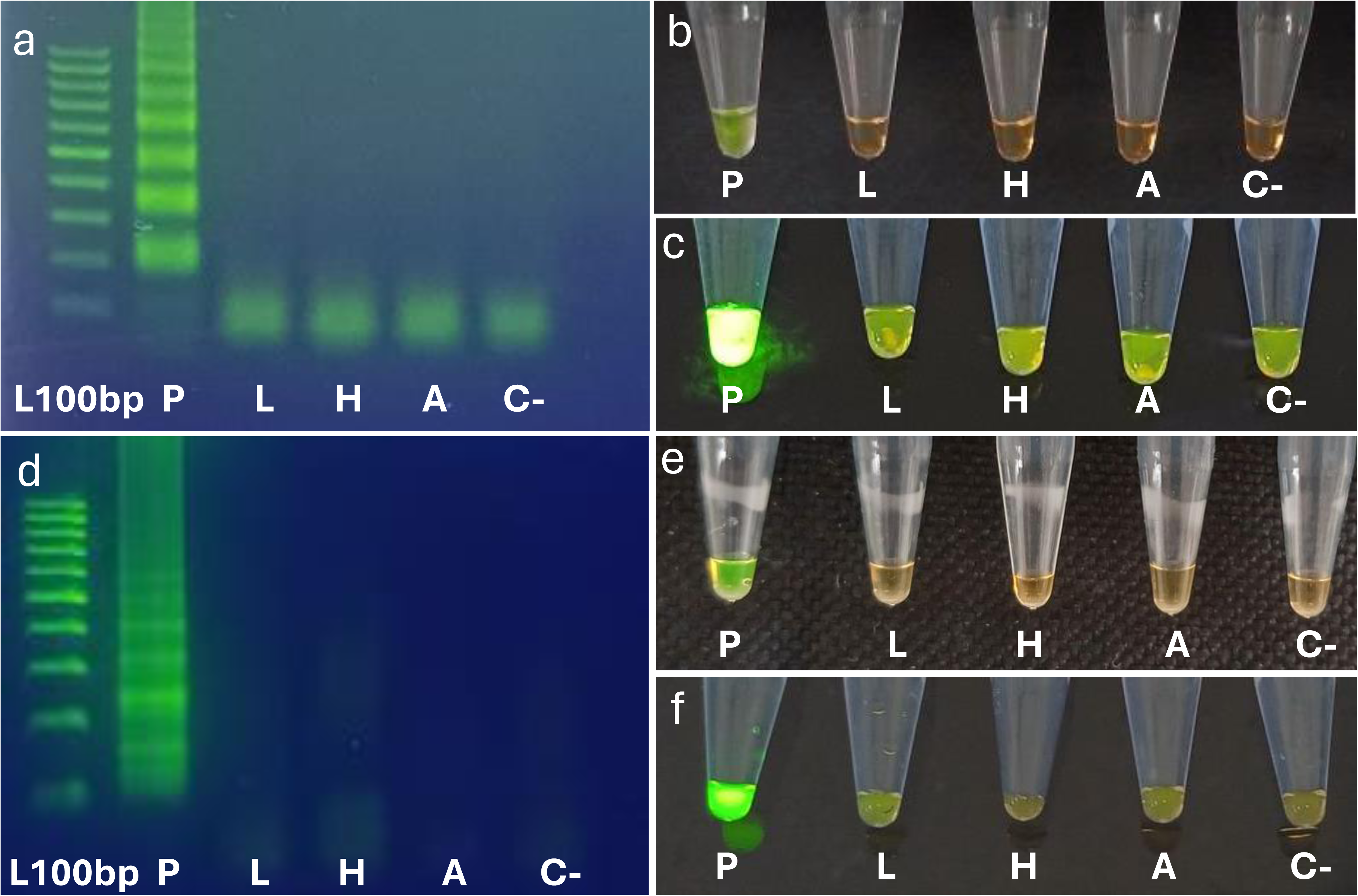
Specificity of LAMP assays. Amplification of *Prorocentrum cordatum* (P), *Prorocentrum lima* (L), *Prorocentrum hoffmannianum* (H), and *Takayama acrotrocha* (A) DNA at concentrations of 10 g/μL, assessed by: (a) and (d) electrophoresis on a 2% agarose gel; (b) and (e) visible color change after staining with 1000× SYBR Green I (Invitrogen S7563); and (c) and (f) fluorescence under UV light. Panels (a) to (c) show results for the ITS-LAMP assay, and panels (d) to (f) for the *psbA*-LAMP assay under the same conditions. Line L100bp: 100 bp molecular marker; lane C-: negative control (water only). Both assays showed 100% specificity for *P. cordatum*, with no amplification observed for the other species.

Consistent with our results, previous studies have successfully applied LAMP and qPCR assays for the specific detection of harmful dinoflagellates, including *Prorocentrum* spp. (Chen *et al*., 2013; McLennan *et al*., 2021; Yang *et al*., 2024; Zhang *et al*., 2014), *Alexandrium* sp. (Wang *et al*., 2008; Zhang *et al*., 2012), *Amphidinium* spp. (Lee *et al*., 2021; Wang *et al*., 2019), *Chattonella* spp. (Qin *et al*., 2019), and *Karenia* spp. (Huang *et al*., 2020; Wang *et al*., 2020; Zhang *et al*., 2009).

Accordingly, the ITS1 and ITS2 regions have previously been used for the specific detection of *P. cordatum* by qPCR (McLennan *et al*., 2021), and, in line with these findings, our ITS-PCR assay also specifically amplified *P. cordatum*. In contrast, this study is the first to demonstrate specific amplification using the *psbA*-PCR assay, as this molecular marker has been comparatively less explored.

### Field sample validation in Conceição lagoon

*Celular detection – Prorocentrum cordatum* was identified in 32 of the 48 field-collected samples by Utermöhl quantification, with occurrences recorded across all six sampling sites (A – F, Figure 1) in Conceição lagoon (Table 2). This dinoflagellate is considered harmful because of its bloom-forming capacity and potential toxicity (Lundholm *et al*., 2009; Heil *et al*., 2005). In South America, a bloom of *P*. *cordatum* was reported in Peru in 2017 (Tenorio *et al*., 2022), while in Brazil cell densities reached up to 1.7 × 10 cells L^−1^ in Paranaguá Bay in 2003 (Mafra Jr. *et al*., 2006). In Santa Catarina, the species has also been recorded in Penha and Florianópolis, with densities up to 7.2 × 10^3^ cells L^−1^ (Alves *et al*., 2010; Miotto and Tamanaha, 2012). Taken together, these findings demonstrate that *P. cordatum* is a recurrent and widespread component of the coastal plankton community in southern Brazil, including the Conceição lagoon. Among all sampling sites, D and F exhibited the highest cell densities in whole (quantitative) samples (samples 25 – 28 and 47 – 48), ranging from >10^2^ to 2.85 × 10^3^ cells L^−1^ (Table 2). The elevated abundance of *P*. *cordatum* at these sites is likely linked to its mixotrophic behaviour and ability to thrive under eutrophic conditions, which characterise the southern region of Conceição lagoon (Silva *et al*., 2017; Stoecker *et al*., 1997; Glibert *et al*., 2008; Heil *et al*., 2005; Khanaychenko *et al*., 2019). Although the observed densities may not pose an immediate ecological risk, they highlight the need for continued monitoring and a better understanding of local phytoplankton dynamics, reinforcing the role of *P*. *cordatum* as a potential sentinel species for eutrophication in the lagoon.

*Molecular detection –* To simplify and improve phytoplankton molecular analysis, this study established a syringe-based biomass concentration method that enabled the detection of *P. cordatum* by both PCR and LAMP assays in concentrated and whole field samples, using volumes as small as 5 mL (Table 2, Supplementary Figure 1). Unlike previous approaches, such as that of Zhang *et al*. (2014), which still required centrifugation to concentrate samples, our method relies on syringe filters, providing a practical and truly point-of-care alternative. Moreover, while Lee *et al*. (2021) applied syringe-based LAMP assays to other dinoflagellates (e.g.: *Ostreopsis* cf. *ovata* and *Amphidinium massartii*), our study is the first to adapt this strategy specifically for *P. cordatum*. In addition, we developed LAMP assays targeting the ITS and *psbA* genes, with detection limits of 10^−1^ ng and 10 ng (10^5^ and 10^6^ cells L^−1^), respectively. Although less sensitive than the LSU-targeting LAMP assay developed by Zhang *et al*. (2014), which achieved a detection limit of 10 ^2^ ng, these thresholds are particularly valuable because *P. cordatum* becomes harmful only at concentrations above 10 cells L^−1^ (Mafra Jr. *et al*., 2024). Consequently, our assays are optimized for bloom-relevant conditions, ensuring rapid, field-applicable detection when early warnings are most critical.

Positive amplicons from the *psbA* and ITS PCR assays in nine samples from Group (i) (11 – 15, 17 – 20, Table 2) were sequenced. NCBI database analysis confirmed >99% similarity to *P. cordatum*. In the phylogenetic tree, *psbA* and ITS sequences from these samples clustered with those from a *P. cordatum* culture isolated from Conceição lagoon, forming a well-supported monophyletic group (Figure 8) and confirming species identity. Collectively, these results demonstrate that PCR reliably detects *P. cordatum* in field samples, consistently confirming its presence across sites A – F (Figure 1) and collection periods (2024 – 2025, Table 2). LAMP assays detected *P. cordatum* in only three samples from Group (i) (4, 14, and 15) using the ITS marker, all with cell densities above 10 cells L^−1^, and none using the *psbA* marker (Table 2, Supplementary Figure 2). This was expected, since ITS assays were more sensitive than *psbA* assays, most likely due to differences in gene copy number (see *Sensitivity* section for details). Also, although the heat-based DNA extraction method appeared slightly less efficient than the commercial kit in Group (i) samples, it processed only 5 mL of sample compared with 50 mL used with the kit, highlighting its practicality for rapid, small-volume, point-of-care applications.

**Figure 8.**
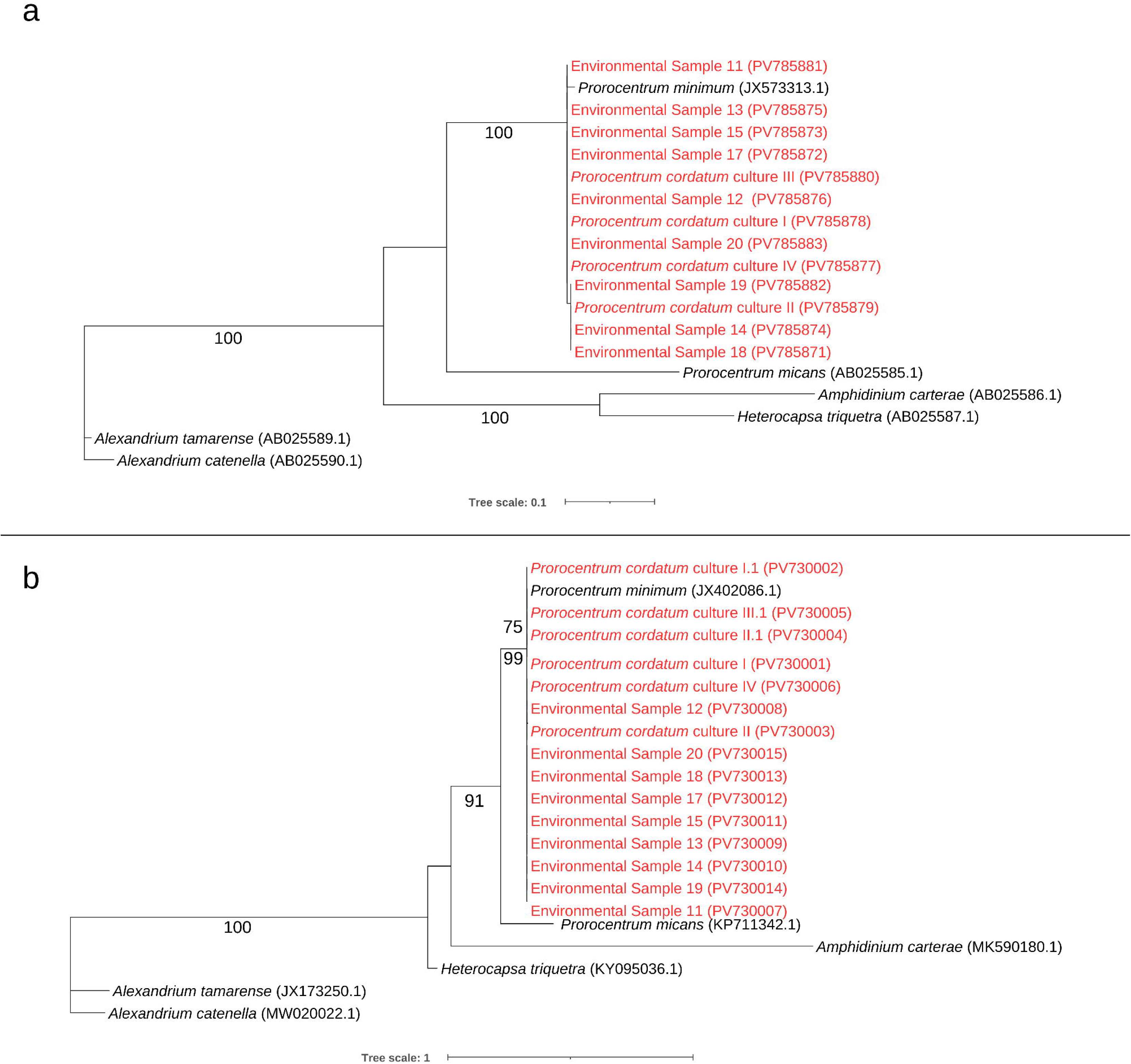
Maximum likelihood *Prorocentrum cordatum* phylogenetic trees based on *psbA* (a) and ITS (b) sequences, constructed using the TVM+F+I and K2P+G4 models, respectively. The analysis included nine sequences from environmental samples (11 – 15 and 17 – 20, Table 2) obtained in this study. Additionally, six ITS and four *psbA* sequences from *P. cordatum* isolates in this study (I – IV) were analyzed, together with additional dinoflagellate sequences used as outgroups (accession numbers in parentheses). The results strongly support that all nine environmental samples belong to *P. cordatum*, as they share monomorphic *psbA* and ITS profiles and form a monophyletic clade with *P. cordatum* isolates from Florianópolis and South Korea (contigs JX573313.1 and JX402086.1). Bootstrap values (based on 1,000 replicates) are shown at the nodes; only values above 75% are displayed.

The simplified protocol combining sample collection, biomass concentration, and heat-based DNA extraction proved effective for detecting *P. cordatum* in both concentrated and whole samples from Group (ii) using PCR. Each sample was analyzed in duplicate using both heat- and kit-based extraction methods, and PCR successfully detected *P. cordatum* in seven of the eight samples with both ITS and *psbA* markers. In contrast, all LAMP assays were negative, consistent with Utermöhl counts of around 10^3^ cells L^−1^, which fall below the LAMP detection limit (>10 cells L^−1^). Together, these findings demonstrate that heat-based DNA extraction combined with whole sampling is a practical and efficient approach for detecting *P. cordatum* in environmental samples with PCR, simplifying both collection and extraction steps and enabling potential field-based analysis.

Samples from Group (iii) (29 – 48, Table 2) were analysed exclusively with heat-based DNA extraction to compare phytoplankton and Van Dorn-bottle collection methods. ITS-PCR detected *P*. *cordatum* in nine samples, while *psbA*-PCR detected it in seven (Table 2, Supplementary Figures 3 – 4). Notably, concentrated sample 33 showed <10^2^ cells L^−1^ in the Utermöhl analysis, yet both PCR assays (ITS and *psbA*) successfully amplified *P*. *cordatum* DNA. A similar result was observed in concentrated samples 41 and 42, where ITS-PCR detected *P*. *cordatum* despite Utermöhl counts of <10^2^ cells L^−1^.

In summary, across all three sample groups (Table 2), PCR assays successfully detected *P. cordatum* in both concentrated and whole samples, with ITS-PCR identifying the species in 32 samples and *psbA*-PCR in 29 (Table 2, Supplementary Figures 3 – 4). PCR also proved more robust and sensitive than microscopy, detecting *P. cordatum* in samples 33 and 41 – 44 from Group (iii), despite Utermöhl counts indicating <10^2^ cells L^−1^. In contrast, LAMP assays returned positive results only in concentrated samples: ITS-LAMP detected *P. cordatum* in three cases (samples 4, 14, and 15), whereas *psbA*-LAMP detected none (Table 2, Supplementary Figure 2). These outcomes are consistent with spiked-sample trials, reflecting the lower sensitivity of *psbA*-LAMP (limit of detection ∼10 cells L^−1^) compared with ITS-LAMP (∼10 cells L^−1^) (Table 2, Figure 5). Although less sensitive than PCR, LAMP represents a practical alternative for monitoring harmful microalgae (Bruce *et al*., 2014; Toldrà *et al*., 2020). Its sensitivity is appropriate for detecting *P. cordatum* at moderate to high cell densities, such as those preceding or during bloom events, when the species becomes harmful (Mafra Jr. *et al*., 2024). In this study, ITS-LAMP reliably detected *P. cordatum* within this range and, when combined with syringe-based biomass concentration and heat-based DNA extraction, provided a simple, field-ready tool. This approach allows detection above baseline levels, providing early warning of bloom development while minimizing dependence on laboratory infrastructure.

This study presents a validated set of molecular methods for detecting *Prorocentrum cordatum*, providing a rapid and broadly applicable tool for phytoplankton monitoring in coastal waters. PCR and LAMP assays proved complementary: PCR offered a highly sensitive and specific laboratory-based method capable of detecting *P. cordatum* at low cell densities, whereas LAMP provided a practical, field-friendly alternative suitable for detecting higher concentrations typical of bloom conditions. Across different sites and cell densities, both approaches confirmed their effectiveness, with the ITS region showing greater sensitivity than the *psbA* gene. In addition, a simple and efficient sampling strategy was established for both concentrated and whole samples. Biomass could be concentrated using syringe filters using volumes as small as 5 mL, while a rapid, low-cost DNA extraction method requiring only nuclease-free water and a heat block proved effective in both field and laboratory settings. Together, these methods deliver a complete and adaptable workflow for detecting harmful dinoflagellates, supporting environmental risk management and enhancing our understanding of *P. cordatum* dynamics in coastal lagoon systems.

## Acknowledgements

The authors are grateful to PDTIS-FIOCRUZ for access to the DNA sequencing facility, LAMEB-UFSC for providing infrastructure and equipment, and Dr. Luiz L. Mafra Jr. from the Universidade Federal do Paraná for donating the algal cultures. We also acknowledge support from FAPESC, CNPq, CAPES, and Rede Brasileira de Pesquisas sobre Mudanças Climáticas Globais (Rede Clima). This paper forms part of the PhD thesis of Andre Akira Gonzaga Yoshikawa, developed within the Programa de Pós-Graduação em Biologia Celular e do Desenvolvimento (PPGBCD) at the Centro de Ciências Biológicas (CCB), Universidade Federal de Santa Catarina (UFSC).

## Author contributions

A.A.G.Y. drafted the original manuscript and participated in all data generation and experimental design. A.A.G.Y, B.V.R., J.V.C.G., S.F.C., P.A.H and L.R.R collected the environmental water samples. A.A.G.Y, B.V.R., J.V.C.G., S.F.C., I.C.P., A.N.P., and L.D.R.P. contributed to data generation and analysis. B.V.R. assisted in cell culture and molecular analysis. S.F.C and J.V.C.G. contributed to primer design, experimental design, and molecular analysis. P.A.H., L.R.R. and C.Y.B.O. supported project logistics, cell culture, quantification analysis, and infrastructure. A.N.P. and L.D.P.R. contributed to drafting by critically reviewing the manuscript. A.N.P. and L.D.P.R. were the principal investigators and participated in the design and coordination of the study. All authors read and approved the final manuscript.

## SUPPLEMENTARY FIGURES

**Supplementary Figure 1.** Biomass concentration by filtration. (a) Syringe (Descarpack) fitted with a syringe filter (Millipore SX0002500 and SX0001300); (b) Field samples pre-filtered using a 100 μm nylon membrane (Millipore, NY1H2500); (c) and (d) Mixed cellulose esters (MCE) 0.22 μm membrane (Millipore GSWP01300) before and after biomass concentration, respectively.

**Supplementary Figure 2.** ITS-LAMP field sample analysis. DNA at concentrations of 10 g/μL assessed by visible colour change after staining with 1000× SYBR Green I (Invitrogen S7563). Line L100bp: 100 bp molecular marker; lane C-: negative control (water only); C+: positive control with *P*. *cordatum* gDNA.

**Supplementary Figure 3.** ITS-PCR (243bp) field sample analysis. DNA at concentrations of 10 ng/μL. Line L100bp: 100 bp molecular marker; line C-: negative control (water, no DNA); C+: positive control with *P*. *cordatum* gDNA.

**Supplementary Figure 4.** *psbA*-PCR (661bp) field sample analysis. DNA at concentrations of 10 ng/μL. Line L100bp: 100 bp molecular marker; line C-: negative control (water, no DNA); C+: positive control with *P*. *cordatum* gDNA.

---

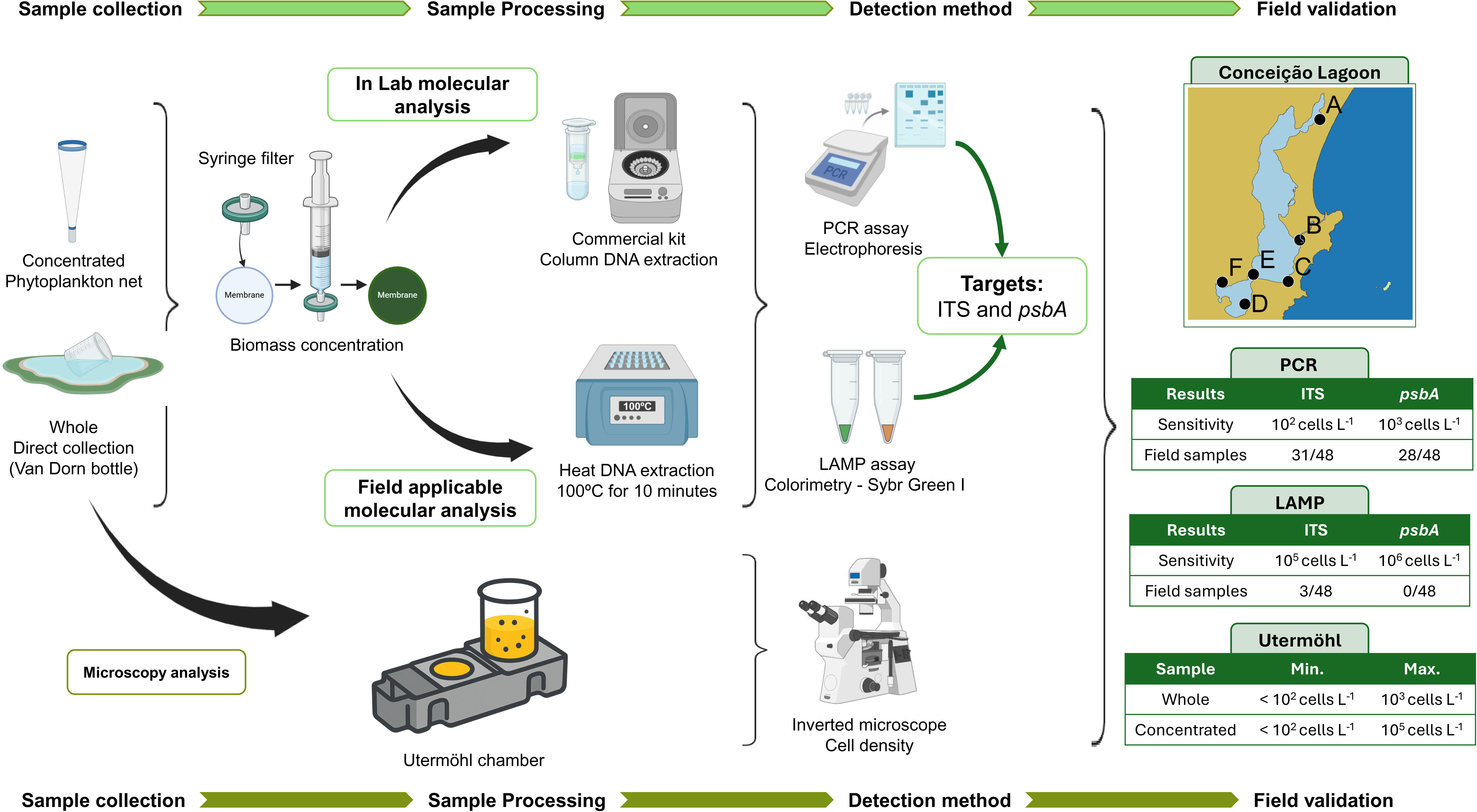

