## Supplementary figures and images for "A field-ready molecular workflow for sample-to-result detection of the harmful dinoflagellate species *Prorocentrum cordatum* in coastal waters"

### Supplementary Figure 1

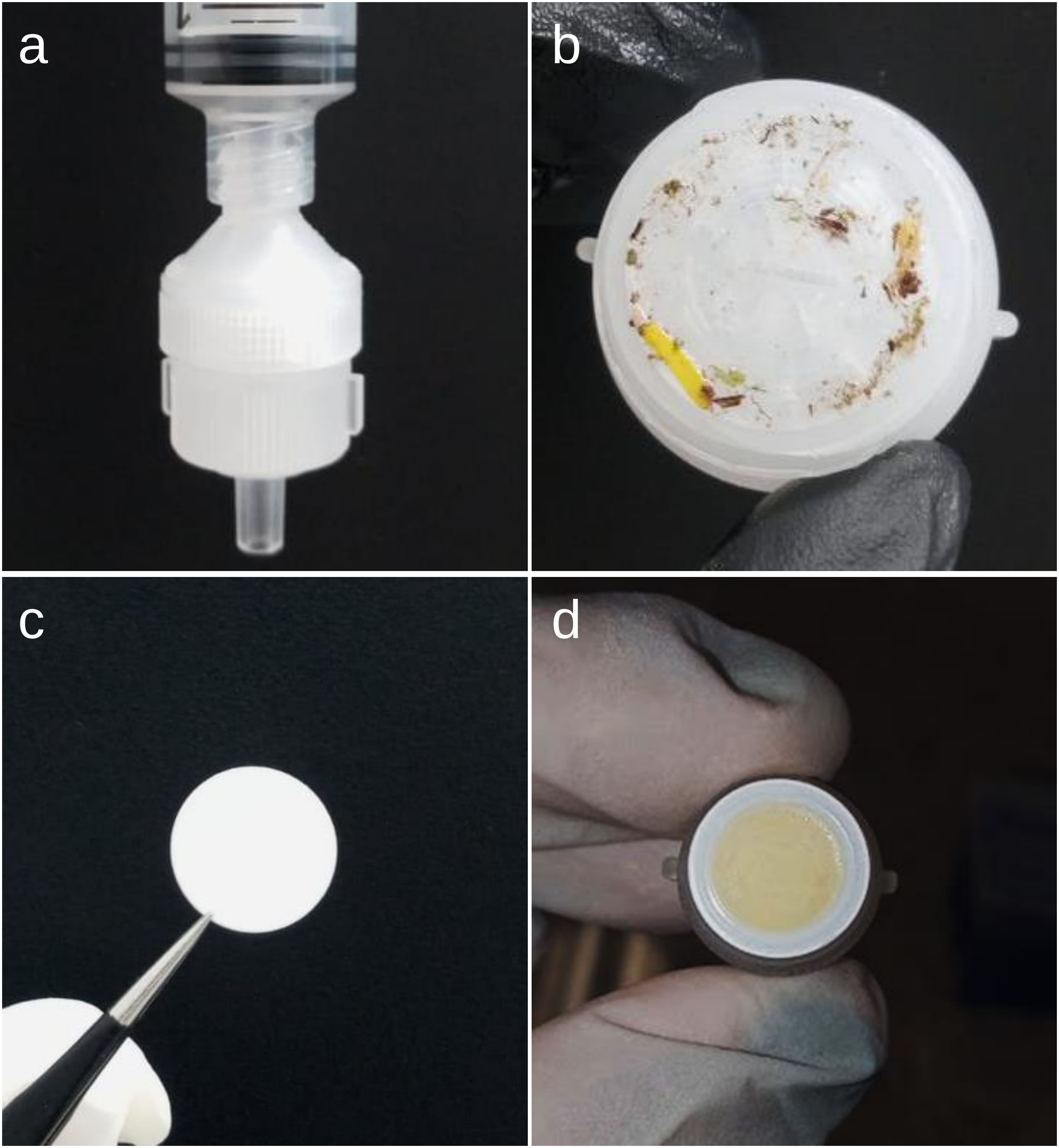

### Supplementary Figure 2

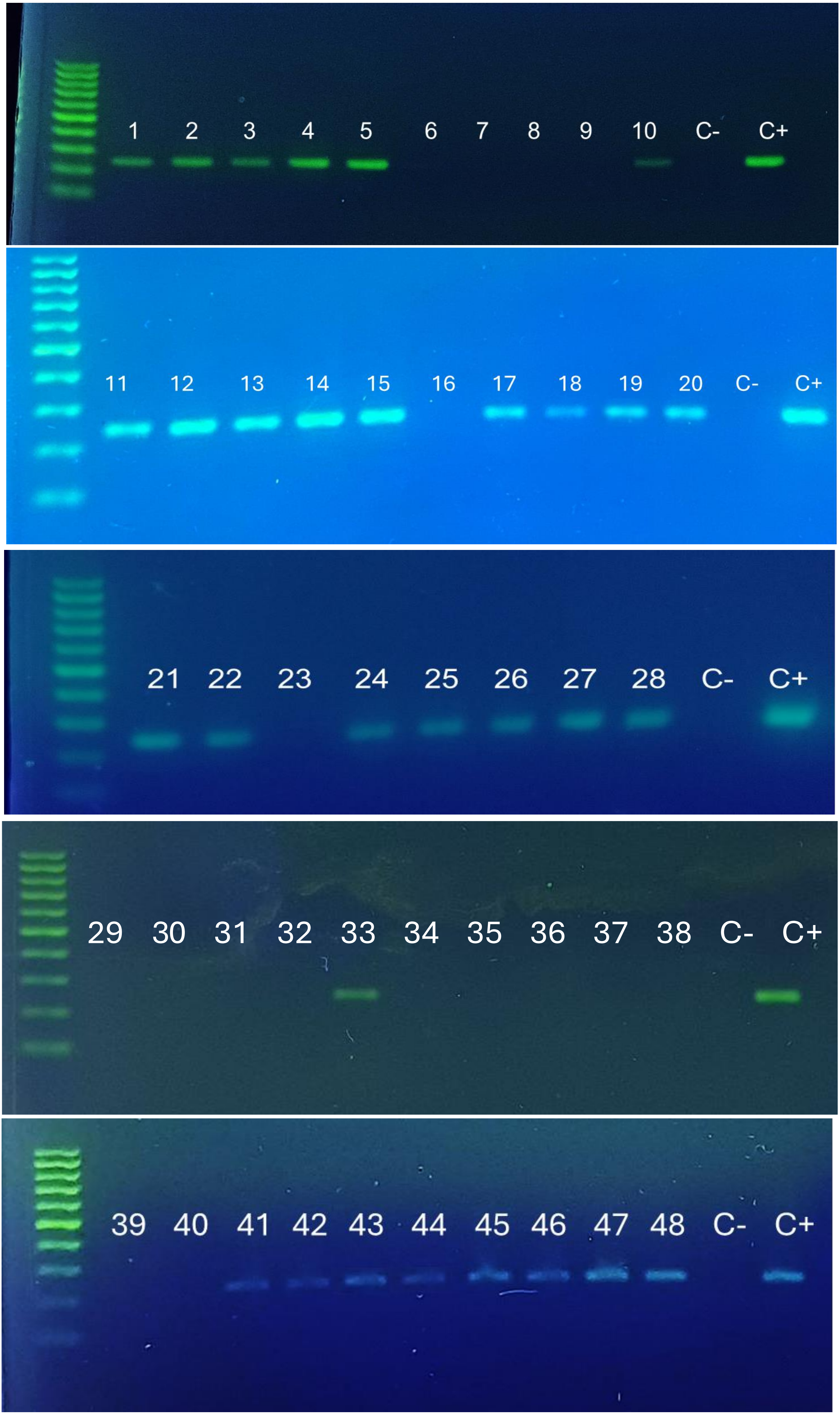

### Supplementary Figure 3

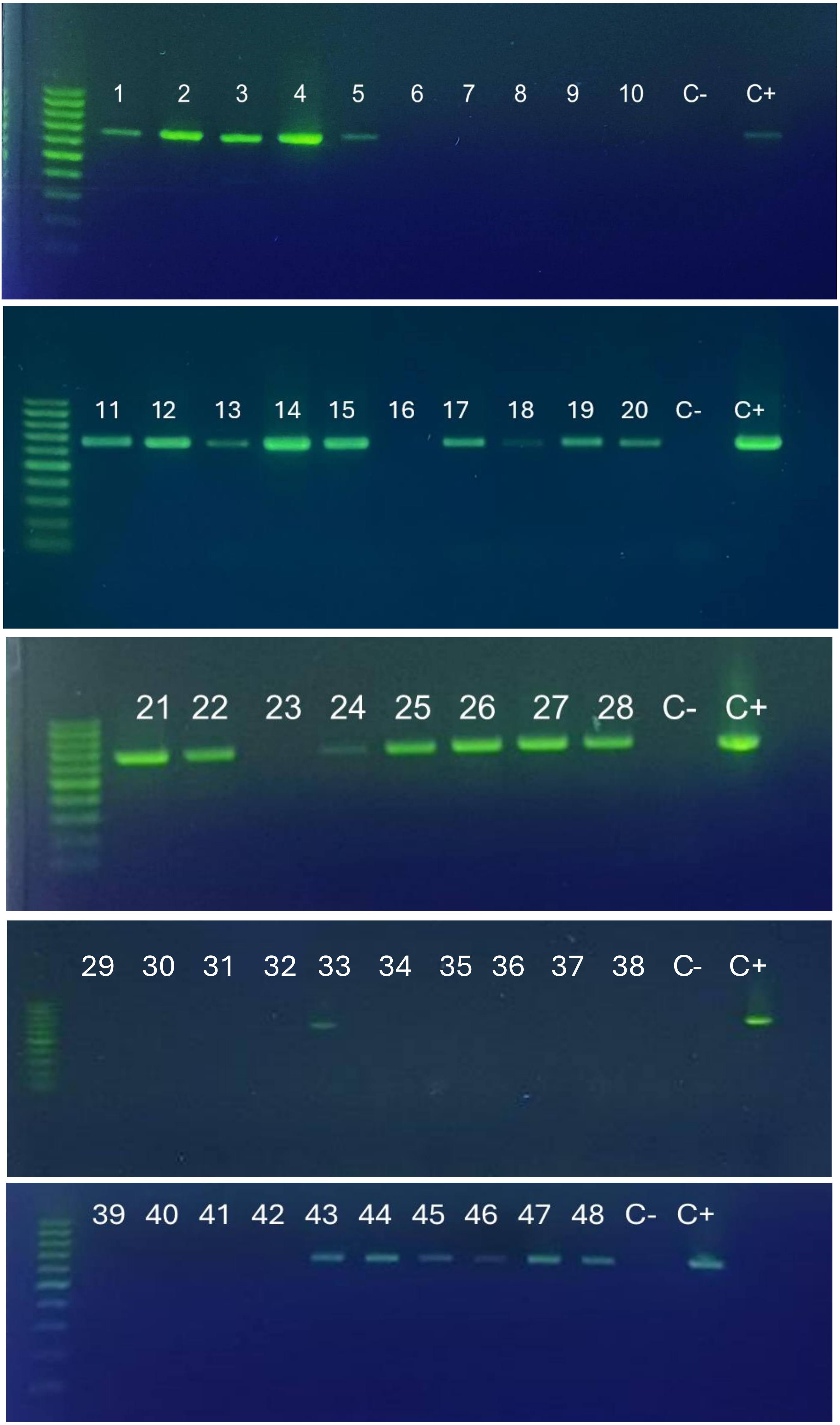

### Supplementary Figure 4

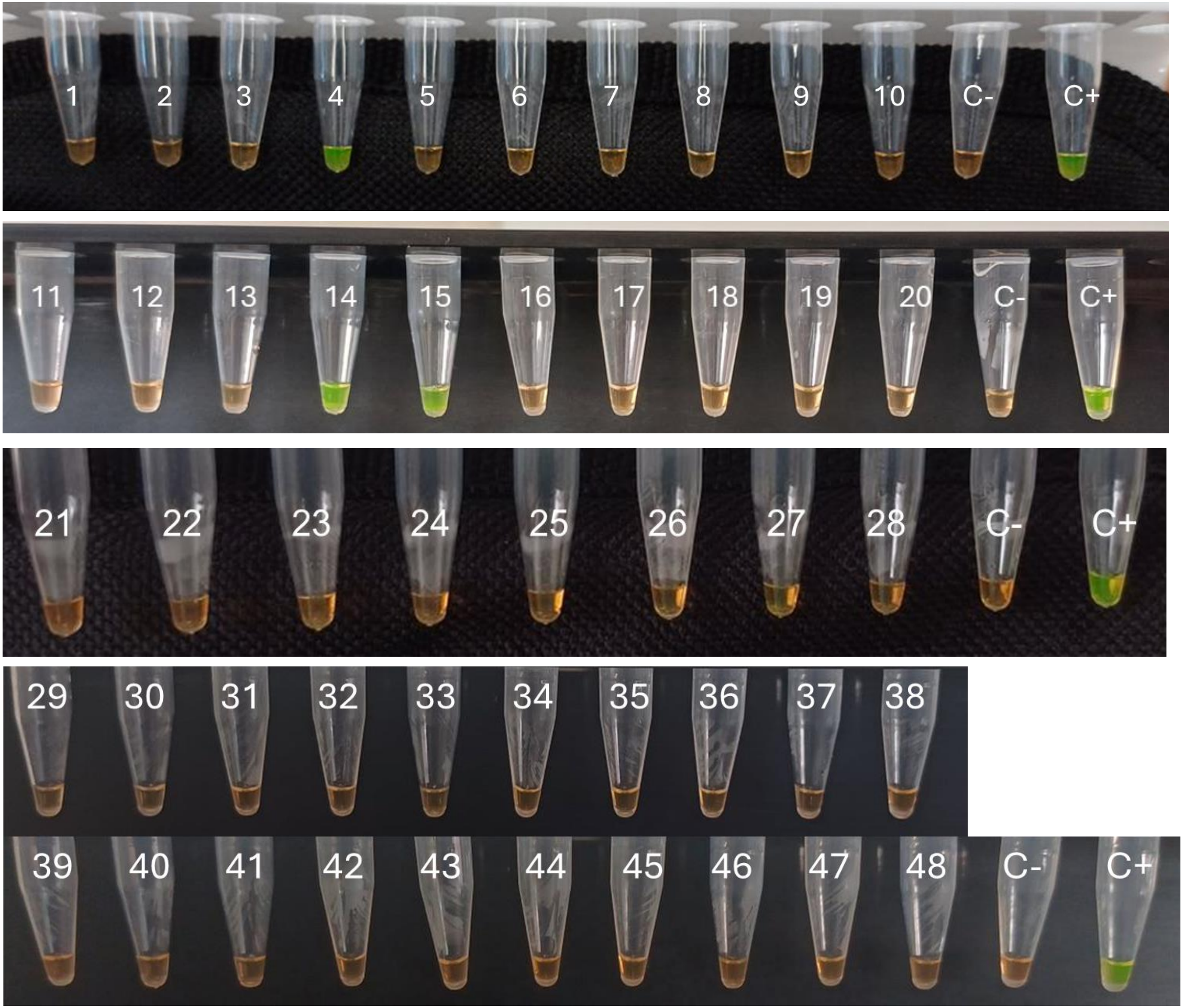
